# Enhancing Strain-level Phage-Host Prediction through Experimentally Validated Negatives and Feature Optimization Strategies

**DOI:** 10.1101/2025.05.31.656987

**Authors:** Min Li, Gufeng Liu, Wenchen Song, Jianqiang Li, Lijia Ma, Minfeng Xiao

**Author notes:** Correspondence (M.X.), (Lijia M.), (Jianqiang L.). These authors contributed equally (Min Li, Gufeng Liu).

## Abstract

**Background:** Accurate prediction of phage-host interactions at the strain level is critical for understanding microbial ecology and for developing phage-based therapeutics. However, existing models are limited by the lack of experimentally validated negative interactions and inconsistencies in data construction strategies.

**Results:** In this study, we present a large-scale phage-host interaction dataset comprising 13,000 experimentally verified links between 125 *Klebsiella pneumoniae((K. pneumoniae)* phages and 104 *K. pneumoniae* strains. Using this unique resource, we systematically evaluate the impact of negative data construction methods, feature extraction strategies, and machine learning algorithms on predictive performance. We show that randomly generated negatives significantly inflate model accuracy, while models trained on experimental negatives yield more realistic and robust results. Furthermore, protein-derived features outperform DNA-based features across various data conditions. Notably, models using only tail protein sequences achieve performance comparable to those using full-genome sequences, offering a time-efficient alternative without compromising accuracy. Finally, interpretable machine learning reveals amino acid preferences in both phages and hosts that align with known infection mechanisms and suggest novel determinants such as anti-transcriptional proteins.

**Conclusions:** Our findings highlight best practices for constructing high-fidelity strain-level phage-host prediction models. The dataset and insights presented here provide a valuable benchmark for future studies and lay the foundation for more biologically grounded, interpretable modeling frameworks in viromics and microbiome research.

## Introduction

Bacteriophage is a class of viruses with host-specificity to bacteria, with numerous amount, diverse morphology, and extensive habitats on Earth[1]. At the beginning of the 19th century, phage was first discovered by *Frederick Twort* and Fe l ix d’He relle, which played a significant role in the microbial dynamics in the ecosystems and the co-evolution with bacteria [2]. With the emergence of multidrug resistant (MDR) bacteria and the challenge of sustaining infection therapy, phage therapy has become one of the most promising alternatives to conventional antibiotics[3], benefiting from a highly specific host range and the minimal disruption of normal flora[4]. In recent years, phage therapy has successfully treated MDR and urinary tract infection in the way of intravenous injection with antibiotics[5, 6], demonstrating the safety and feasibility of phage therapy.

However, there is still a considerable challenge in the clinical application of phage therapy about determining the relationship of phage-bacteria at strain level accurately and quickly[5-7]. Several experimental methods were developed to confirm the relationship and measure the intense of interaction between phages and bacteria, such as spot assay[8], microfluidic-PCR[9] and PhageFish[10]. Although experiment methods are the gold standard of phage therapy, it usually consumes at least several days depending on the host amounts, largely limiting the application of phage therapy [11, 12]. In addition, a major obstacle was the scarce host information of phage genome in public databases, such as NCBI GeneBank, EMBL-EBI, Phantom, and MVP (a microbe–phage interaction database)[13], some of which identified isolates host to only the genus level, posing inconvenience to the phage therapy and phage-host research. Therefore, improving the precise matching of phage and host at strain level was necessary and valuable.

Computational methods can serve as a complementary or pre-experimental approach to help researchers efficiently narrow down experiment scale. Currently, many tools for phage host prediction have been developed, categorized into two types based on alignment-dependent or alignment-free. Alignment-dependent phage-host prediction tools difficultly identify hosts in the absence of related species or related phage-host pairs due to relying on sequence similarity, demanding a high level of comprehensiveness from the reference database. Alignment-free tools extracted the contigs characteristics like k-mer frequency, properties of protein sequences, and PseDNC as features to fit custom formulas[14], Homogeneous Markov model[15], Gaussian[16], Logistic matrix factorization[17], similarity network fusion[17], or train machine learning[18], deep learning model[19], reducing the dependence of sequence similarity. Most phage host prediction tools could be helpful to narrow down the genus or species level to screen (for example, HostPhinder[17]). Now available strain-level phage-host prediction tools include Leite’s method[20], PrePHI[21], and PHIAF[19], which gathered phage-host interaction pairs from publicly accessible databases and generated non-interaction pairs through randomly selecting, K-Means clustering, and GAN augmentation to construct a balanced training set. Leite’s method and PredPhi extracted features by calculating amino acid frequency, molecular weight, and chemical composition based on all protein-coding sequences. PHIAF fused the features originating from all DNA and protein sequences.

The data deficit was a tremendous defect for constructing a precise strain-level phage-host prediction framework, given that the negative infection information is unavailable in the public database and the host information is mostly restricted to the native host. Due to missing experimental negative infection data, the validity of the negative-generation method adopted by Leite’s method, PrePHI, and PHIAF had not been evaluated, and the prediction performance bias from the manual dataset has never been estimated. Additionally, the complex biological process of phage-host interaction involves some key protein interactions[22, 23], such as bacteriophage receptor binding proteins. Tail proteins are a type of receptor binding protein and play a core role in phage-bacteria interaction, which can specifically recognize and bind to capsular polysaccharides and antigens on the surface of bacteria. However, the contribution of genome, proteins and specific protein to machine learning models has not been systematically characterized. Finally, interpretability was an essential aspect of biological research, while the complexity of ML models made them challenging to interpret[24].

In this study, we focused on evaluating the construction method of negative pairs and precise phage-host interaction prediction strategy establishment at strain level for *K. pneumoniae,* which is one of the common multi-drug resistant strains in hospital. The large phage-host experimental dataset was collected from CNSA(Methods ‘’), consisting of 12, 062 negative and 938 positive links about 104 *K. pneumoniae* strains and 125 *K. pneumoniae* phages. Based on the dataset, we investigated the structural differences and modeling effectiveness between the proposed negative data generation approach (random-selecting) and experiment dataset, feature strategy, and model interpretability to reveal the biological mechanism(Figure 1).

**Figure 1.**
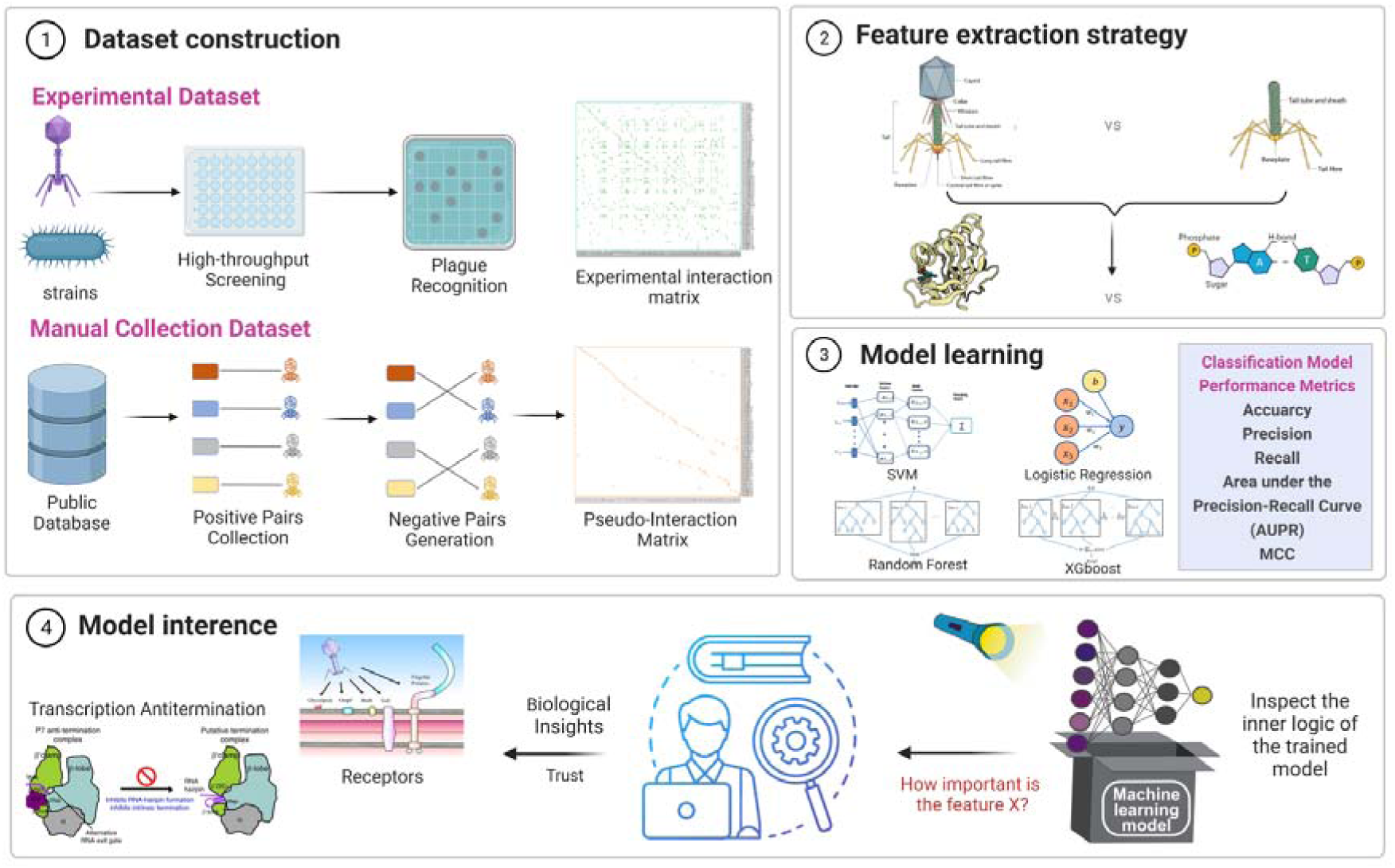
Framework of enhancing phage-host prediction at the strain-level resolution. (1) Dataset Construction Workflow: Illustrates the design and characteristics of the experimental dataset and the pseudo-infection dataset. (2) Feature Extraction Strategy: Incorporates features derived from wholegenome sequences, tail sequences, as well as protein and DNA-level characteristics. (3) Model Training and Evaluation: Support Vector Machine (SVM), Logistic Regression, Random Forest, and XGBoost algorithms are employed to train the models and evaluate their performance. (4) Biological Insights: Key biological proteins and potential infection mechanisms are inferred from model interpretation.

Our findings indicate that models trained on experimental negatives outperform those using randomly generated data, avoiding inflated performance metrics and improving robustness. Protein-derived features, particularly those extracted from phage tail proteins, were found to be more predictive than DNA-based features, achieving comparable accuracy to full-genome models with significantly reduced computational cost. Moreover, interpretable models revealed key amino acid signatures associated with host recognition, providing novel insights into phage-host interaction mechanisms. Collectively, our study offers a benchmark framework and best-practice recommendations for accurate and interpretable phage-host prediction at the strain level.

## Results

### Comparison of network structure between experimental and manual dataset

Phage-bacteria infection network provides a unifying lens to unveil commonalities of microbial and viral communities[25]. Ecological network structure and statistical analysis were carried out based on the strain-level phage-host experimental dataset collected from CNSA. The results showed that the experimental dataset exhibited obvious modularity with statistically significant (Q=0.6, p-value<0.05) and significant nestedness (NODF = 9.98, p-value < 0.05), which is consistent with the phenomenon of phage-host interaction communities in previous studies [25](Figure 2). In addition, the average number of phages that infected a given host was 7.5 ± 6.2, while the average number of hosts that phages could infect was 9.01 ± 8.61 (Table 1).

**Figure 2.**
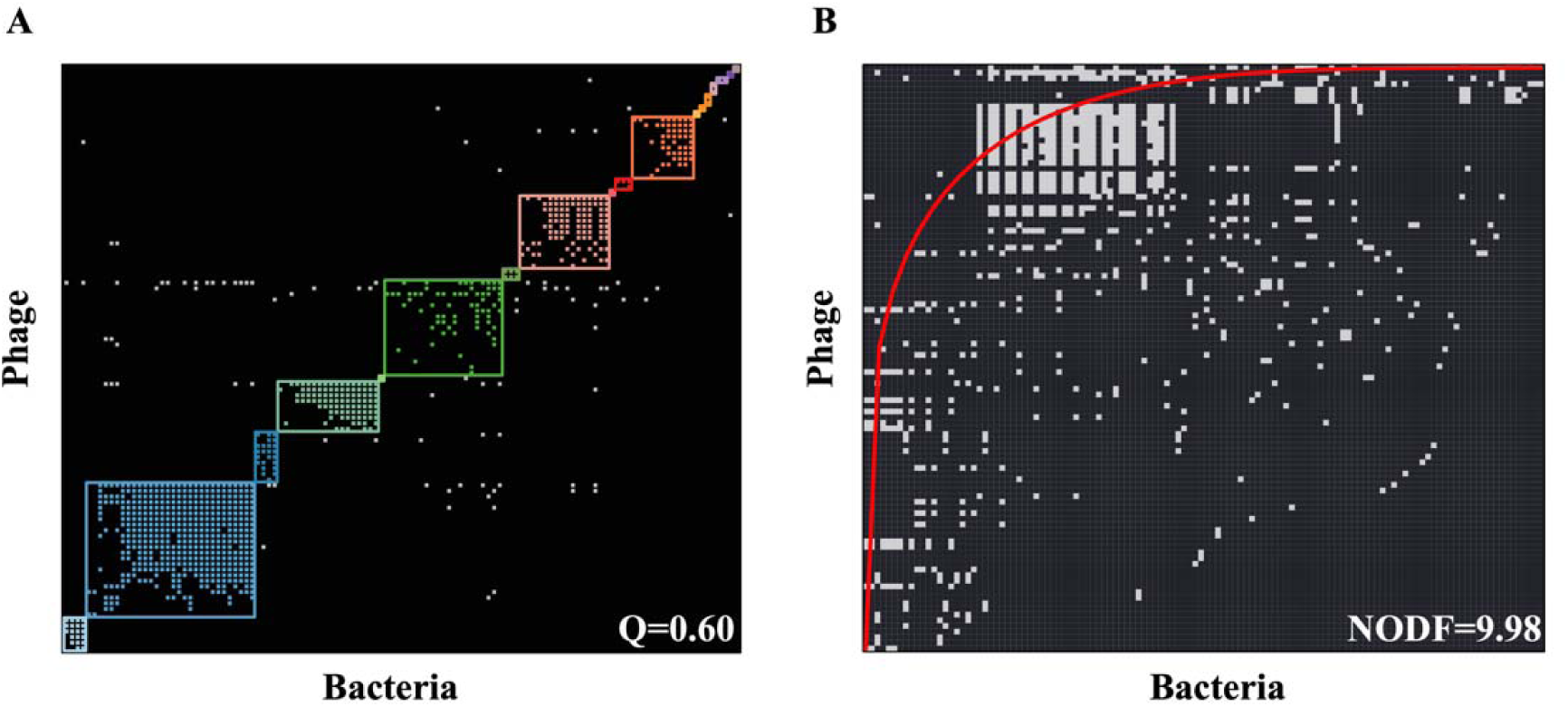
Modularity and Nestedness of the Experimental Phage–Host Interaction Network. (A) Modularity Structure: The modular organization of the experimental phage– host interaction network yields a modularity index of Q = 0.60. Distinct modules are delineated by different colors and borders. White dots located outside the modular blocks indicate positive interactions that are not assigned to any specific module. (B) Nestedness Pattern: The nested arrangement of the same network exhibits a nestedness index (NODF = 9.98). The red curve represents the fitted trend line. Black dots correspond to non-infective phage–bacterium pairs, while white dots denote infective interactions.

**Table 1.** General Characteristics of the Phage–Bacteria Interaction Network.

|  |  |  |
| --- | --- | --- |
| **General properties** | # Host strains | 125 |
|  | # Phages | 104 |
|  | # Interactions (I) | 938 |
| | Size ($M=H \times P$) | 13000 |
| | Connectance ($C = I/M$) | 0.0721 |
| **Hosts** | Infected strains | 120 |
| | Mean (±sd) phage infection per strain | $7.5 \pm 6.2$ |
|  | Maximal phage infections per strain | 23(KP8924) |
| **Phages** | Maximal strains infection per phage | 36(PH-KP7569) |
| | Mean (±sd) strain infections per phage | $9.01 \pm 8.61$ |

To make the comparison fair and thorough, a manual dataset was constructed based on the experimental dataset according to the description of Leite’s method, called a randomly-selecting dataset. For individual phages, the native host was used as the reference for positive infection, and other hosts were marked as negative. The hosts showing fully resistance to any phage (rows in the matrix with only 0) were discarded, resulting in a randomly-selecting dataset with 104 phages and 80 bacteria. Compared with the experimental dataset, the randomly-selecting dataset is characterized by the one-to-one infection relationship between phage and bacteria(Figure2). In the experimental dataset, phages could infect a unique host or several phylogenetically adjacent hosts, resulting in nearly diagonal or block matrices that exhibit a high degree of modularity rather than one-to-one. The negative pairs were pointed out as the main difference between the randomly-selecting dataset and the experimental dataset (Figure 3).

**Figure 3.**
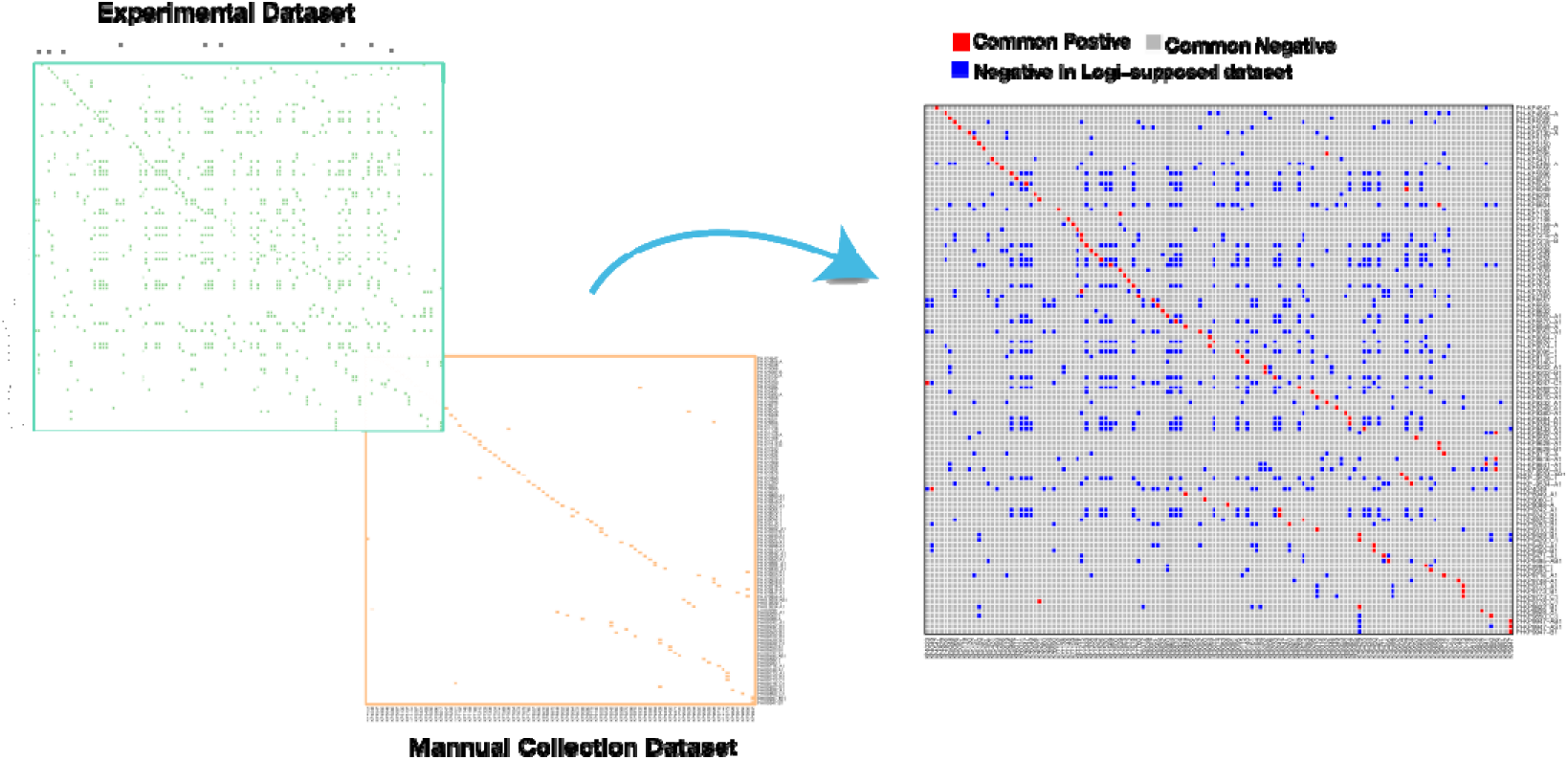
Structural Comparison between the Randomly Selected Dataset and the Experimental Dataset. Legend: The right side of the figure showed the structure of the randomly-selecting dataset and experimental dataset, in which orange and green represent infection, and white represented non-infection; the right side of the figure showed the commonalities and differences between the datasets, in which red represented common positive infections, grey represents common negative infections, and blue represents negative data only in the randomly-selecting dataset. The horizontal axis was the bacteria strain ID, and the vertical axis was the phage ID.

Our results evidence that interaction network structure patterns are different in the experimental and random-selecting datasets, indicating the importance of true negative pairs in the real-world biology community. The one-to-one pattern in the random-selecting dataset might be caused by the incomplete knowledge of biological mechanism, which cannot provide the full information to machine learning models and might lead to misunderstanding.

### Effectiveness of modeling performance between experimental and manual dataset

To evaluate the effectiveness of two datasets for predictive modeling, we firstly trained several machine learning models based on the experimental dataset with 10-fold cross-validation, such as logistic regression, random forest, SVM, and XGboost, to accurately predict phage-host interaction as the standard of benchmark. As shown in Table 2, XGboost had the best performance in the accuracy, recall, F1, area under the precision-recall (AUPR), and Matthews correlation coefficient (MCC). Therefore, XGboost was used for subsequent performance comparison within the experimental and random-selecting dataset.

**Table 2.**
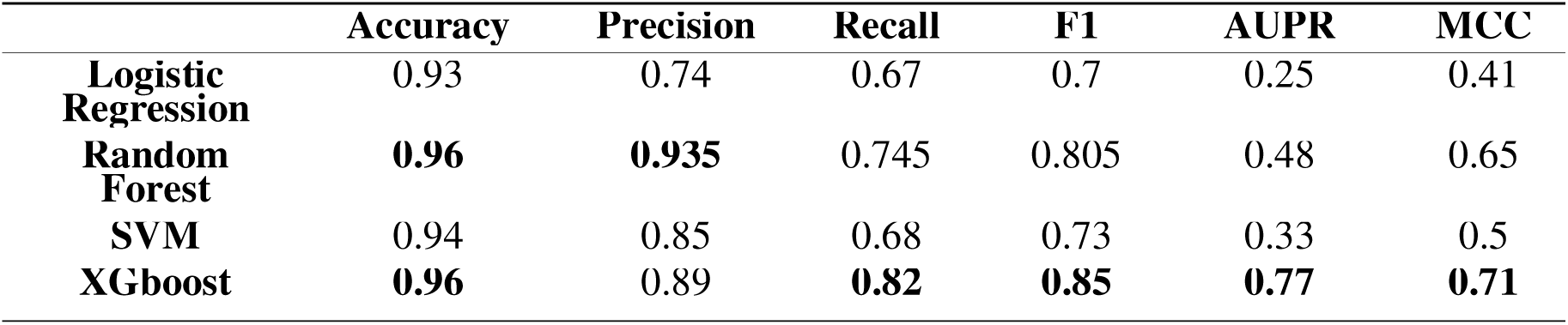
Performance Comparison of Machine Learning Models Trained on Experimental vs. Random-Selection Datasets.

The proportion of positive pairs in the experimental dataset was 7.2% due to the unbalanced characteristics. While Leite’s method evaluated performance of the prediction model through a balanced dataset, we offer fair comparison with the experimental and random-selecting dataset in different positive and negative gradients from 1:1-1:11 based on XGboost(Figure 4). The result shows that with increasing of positive-negative pairs gradient, the accuracy of models trained with experimental and random-selection datasets both improve (Figure 4A). The accuracy of model trained with the experimental dataset was 81%~96% and that of the random-selecting dataset was 52%~92% (Figure 4A; Supplementary Table 1). Generally, the imbalanced dataset has a significant impact on the performance of the model, while the accuracy increases in our result with the imbalanced gradients increasing.

**Figure 4.**
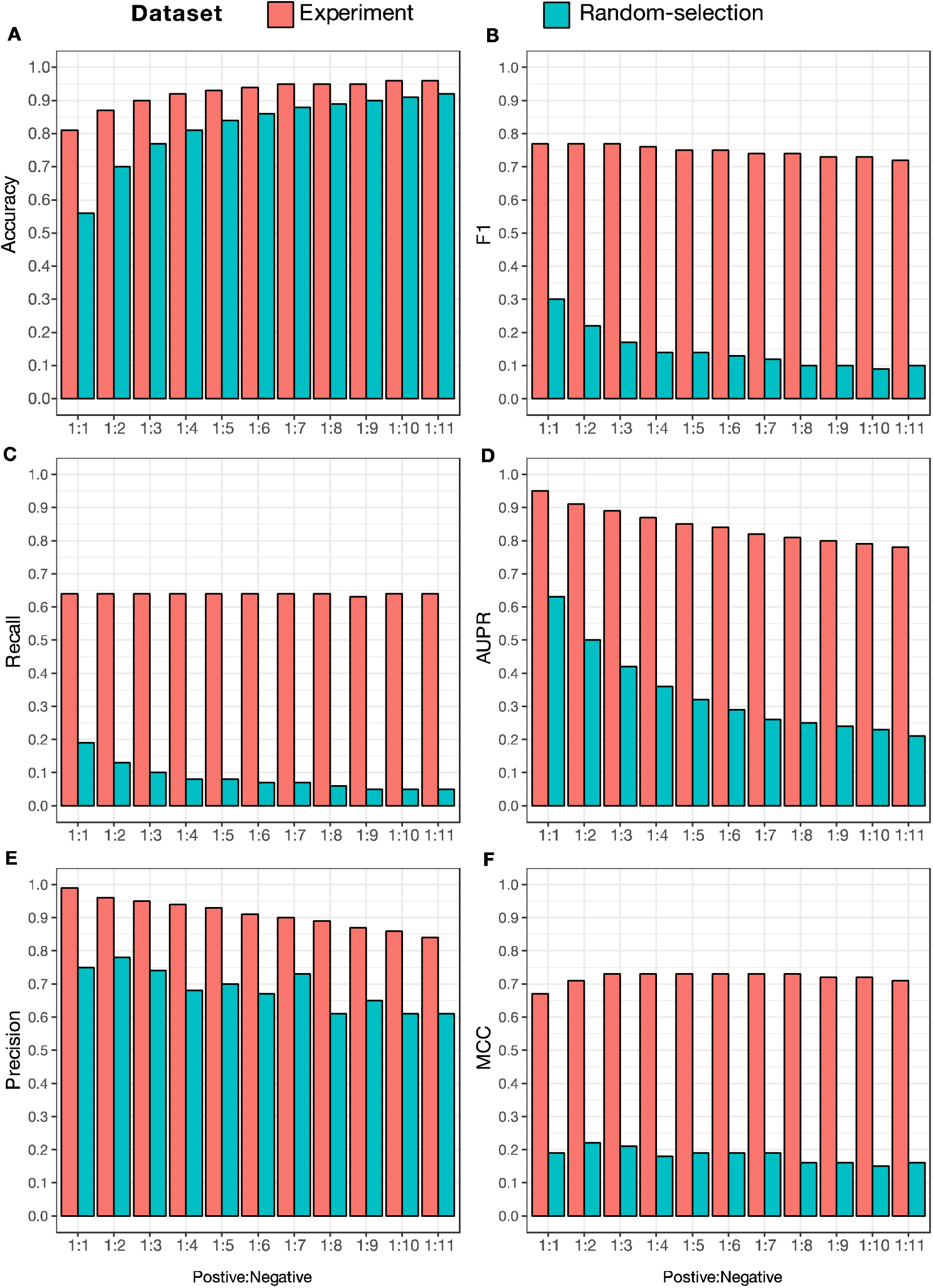
Effectiveness of Experimental and Random-Select Datasets on Model Performance. Comparison of model performance between the experimental group (red) and the random-selection group (blue) under different Positive:Negative sample ratios, including (A) Accuracy, (B) F1 Score, (C) Recall, (D) AUPR (Area Under the Precision–Recall Curve), (E) Precision, and (F) MCC (Matthews Correlation Coefficient).

To prevent the bias of evaluation caused by unbalanced characteristics, we also analyzed the prediction number of true positive and negative. The result shows that the predictive number of true positives decreased in the random-selecting dataset, while more true positive pairs were predicted in the experimental dataset (Supplementary Figure 1). As for the predictive amount of true negative pairs, it increased with the positive-negative ratio increasing in both datasets. Given the imbalanced nature of the datasets, the binary classification model tended to classify the sample as negative, contributing to the true negatives increasing sharply.

In the case of imbalanced data, other unbiased metrics not only the accuracy should also be applied to evaluate the model’s performance, such as precision, recall, the Matthews correlation coefficient (MCC), F1 score and AUPR. The recall, F1 and MCC of the experimental dataset was nearly stable(Figure 4B, 4C, 4F) and that of the random-selecting dataset decreased about 19%∼5% across different positive and negative gradients from 1:1-1:11(Figure 4C; Supplementary Table 1). Precision and APUR in both datasets showed a downward trend(Figure 4E, 4D). Specifically, the precision for the experimental dataset at predicting outcome was 99%~84% and that of the random-selecting dataset was 75%∼61%. The AUPR for the experimental dataset at predicting outcome was 95%∼78% and that of the random-selecting dataset was 63%∼21% (Figure 4D; Supplementary Table 1). Overall, the metrics of the experimental dataset achieved an average of 10% accuracy, 55% recall, 23% precision, and 51% AUPR gain against the random-selecting dataset, respectively (Supplementary Table 1).

To further explore the improvement gained by the experimental dataset, we designed two sets of experiments based on the main difference between the two datasets. One group was composed by the entire random-selecting dataset, and the other used the combination of the positives in the random-selecting dataset and the negatives in the experimental dataset. Compared to completely using the randomly-selecting dataset, the group that added experimental negative data improved an average of 24.8% accuracy, 19% F1, 16.8%% recall, 21.8% precision, and 16.6%APUR (Figure 5A, 5B, 5C, 5D, 5E; Supplementary Table 2). The result indicated that experimental negative pairs played critical role in the model’s performance, significantly increasing the number of true positive samples recognized by the model(Supplementary Figure 2; Supplementary Table 2) and improving the metrics such as accuracy, recall, precision, and AUPR.

**Figure 5.**
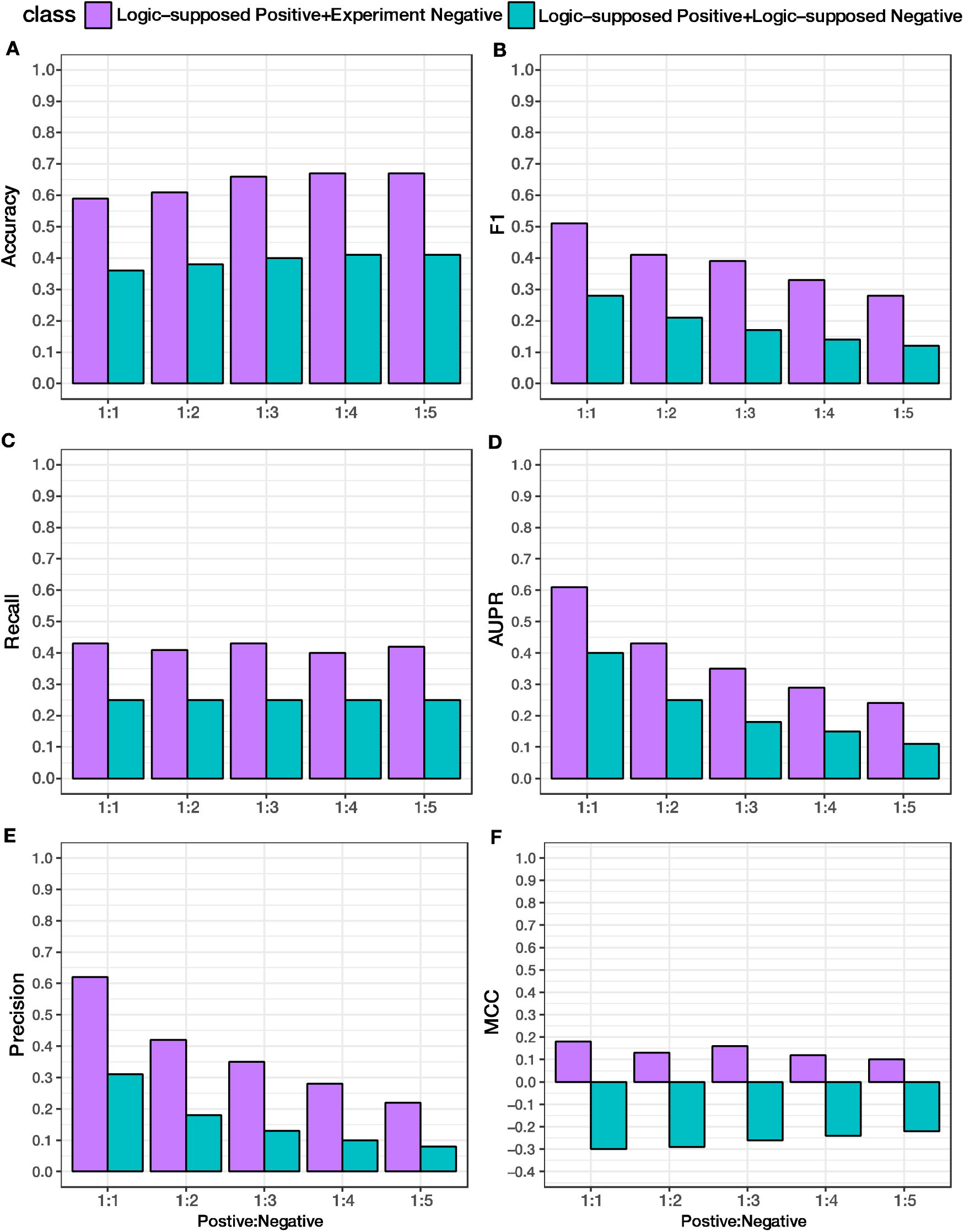
Effectiveness of Experimentally Derived Negative Data on Model Performance. Comparison of model prediction performance metrics between "Logic - supposed Positive + Experiment Negative" (purple) and "Logic - supposed Positive + Logic - supposed Negative" (teal) groups under different Positive:Negative sample ratios, including (A) Accuracy, (B) F1 Score, (C) Recall, (D) AUPR, (E) Precision, and (F) MCC.

### Feature engineering strategy for improving phage-host prediction model

Whole-genome and proteome usually contain a great deal of information, which might overwhelm key information in the interaction between the phage and host, increasing the computation time and harming the robustness of models. It was reported that tail proteins and lysins were strongly associated with the successful infection of phage-host, playing an important role in host recognition and determined the host range with stronger host specificity. However, the effectiveness of tail proteins, whole-genome and whole-proteome in modeling has not been systematically investigated. Considering the protein and DNA sequence features were frequently used in previous studies[15, 19, 26-29], the prediction performance of tail proteins and whole-genome sequences based on DNA and protein features was compared on the experimental dataset to better establish the feature engineering strategy.

In terms of genome sequences, protein features outperformed than DNA features with an average improvement of 1.6% accuracy, 8% recall, 2.3% precision, and 4% APR (Figure 6; Supplementary Table 3). As for tail sequences, protein features improved on average by 1.1% accuracy, 4.9% recall, 0.8% precision, and 2.6% APR than DNA features (Figure 6; Supplementary Table 3). Whether using whole-genome sequences or tail sequences, protein features were superior to DNA features.

**Figure 6.**
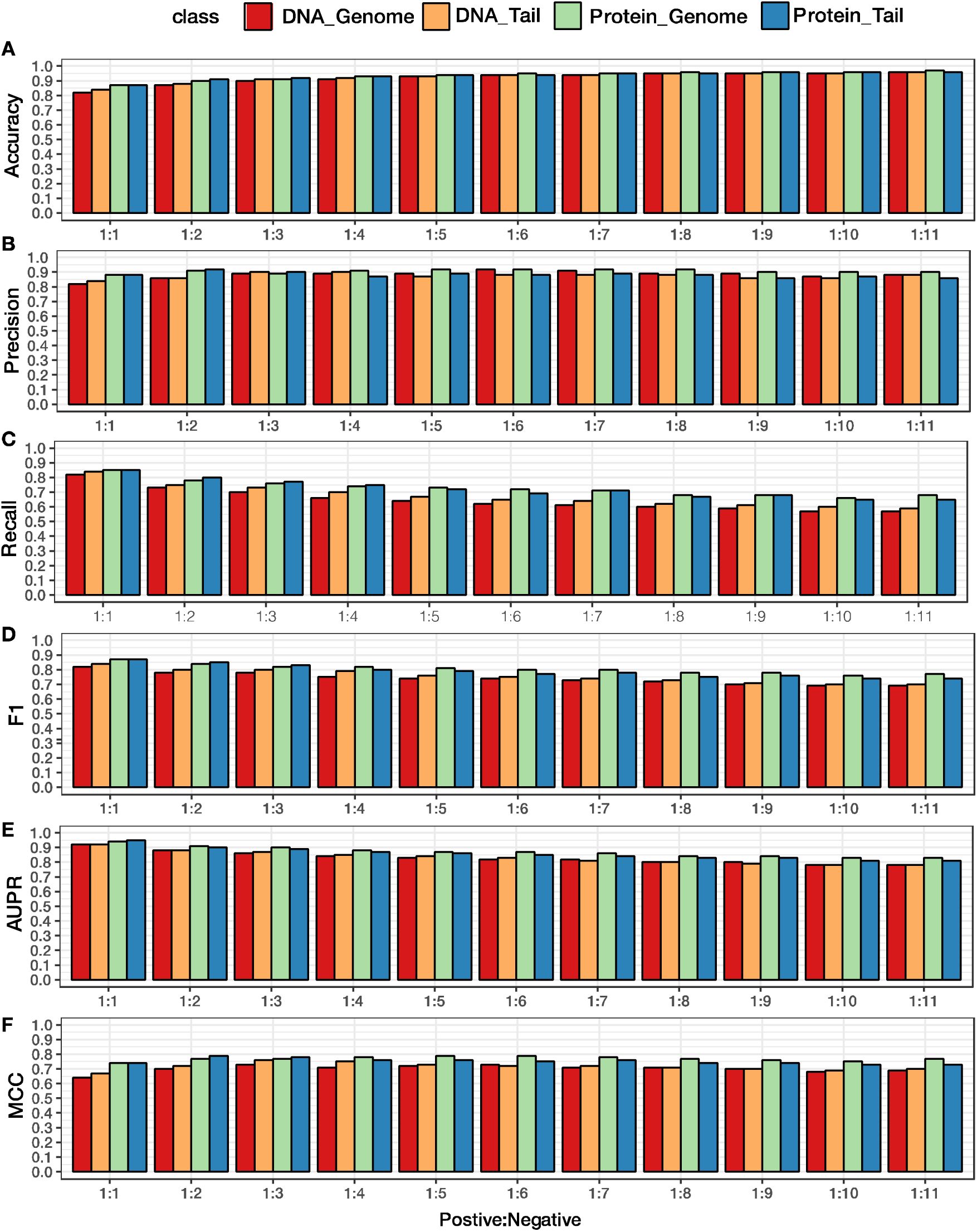
Comparison of Tail vs. Whole-Genome Sequences, DNA Features, and Protein Features in Model Performance. Model performance metrics are compared across four feature combinations such as DNA_Genome (red), DNA_Tail (orange), Protein_Genome (green bars), and Protein_Tail (blue) under different Positive:Negative sample ratios, including (A) Accuracy, (B) F1 Score, (C) Recall, (D) AUPR, (E) Precision, and (F) MCC.

**Figure 7.**
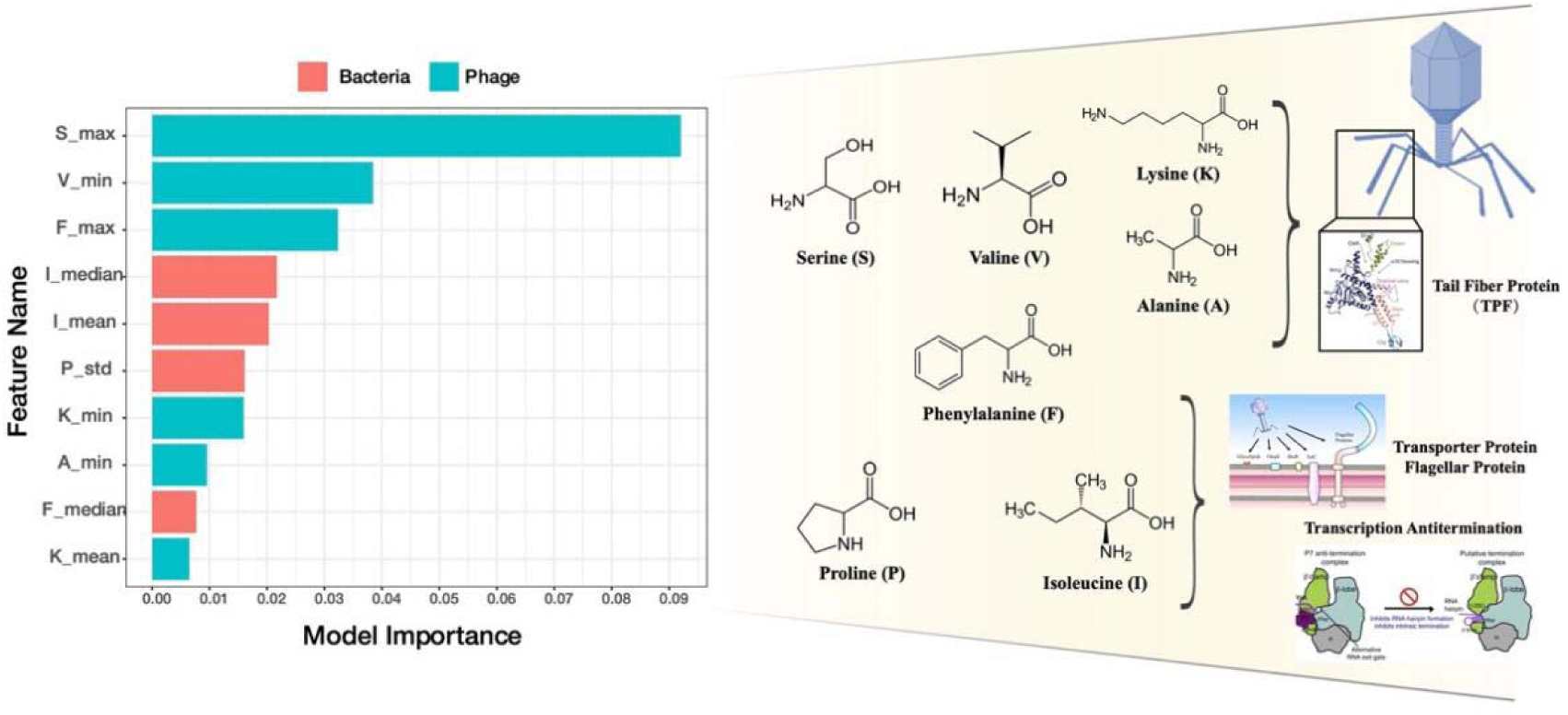
Top 10 Important Features in the Tail Protein-Based Prediction Model. (Left) Bar plot displaying the top 10 most important features contributing to the tail protein model’s classification between bacterial (red bars) and phage (teal bars) proteins. Features are ranked based on their relative importance, with longer bars indicating greater influence on the model’s predictive performance. (Right) Chemical structures of key amino acids, including Serine (S), Valine (V), and Lysine (K), which are likely biologically relevant in distinguishing phagehost interaction. Diagrams illustrating a transporter protein and a transcription anti-termination protein that contains a high abundance of key amino acids, representing potential biological pathways or molecular mechanisms influenced by the identified predictive features.

On the other hand, tail sequences demonstrated performance comparable to that of whole-genome sequences when protein features were utilized. Specifically, the combination of tail sequences and protein features exhibited only marginal differences compared to the whole-genome and protein feature combination, with average gaps of 0.09% in accuracy, 2.45% in precision, 0.45% in recall, and 1.18% in AUPR. Notably, when the imbalance ratio between positive and negative samples was less than 1:4, tail sequences even outperformed whole-genome sequences in terms of precision, recall, and accuracy.

We systematically evaluated the impact of different data sources, including tail and whole-genome sequences and feature characterization methods, such as protein-based and DNA-based features on strain-level phage-host prediction modeling. The results indicate that protein features consistently outperform DNA features across various positive-to-negative ratios. Moreover, tail protein sequences achieve performance comparable to that of whole-proteome sequences.

### Model Interpretability reveals Biological mechanisms

Machine learning (ML) able to find complex patterns in high dimensional and heterogeneous data has emerged as a powerful tool in bioinformatics. But the complex structure underlying ML models made it hard to understand for humans, commonly called as a black box. Recently, advances in the field of interpretable ML have made it possible to identify important patterns and features underlying an ML model. These interpretation strategies can be applied in genetics and genomics to derive novel biological insights from ML models.

In this study, we presented that the important features in the XGboost model for predicting phage-host interaction were utilized to reveal the underlying factor in the phage-host interaction mechanism. The predictive weight of each feature was obtained from the XGBoost model trained with the tail protein sequence and protein property, which showed that the S,V,F,K,A amino acid in phage and the I,P,F amino acid in bacteria had larger weights thus contribute more to the model predicted outcome. Then, the essential proteins were deduced from the amino acid frequency, respectively phage tail fiber proteins with higher amino acids frequency (S, V, F, K, A) and for bacteria, the transporters, fimbriae chaperones, anti-transcriptional proteins with the highest amino acids frequency (I, P, F). Previous studies have experimentally shown that the phage tail fibers play an essential role in phage adsorption and binding to the bacterial cell receptor, such as transporters, and fimbriae chaperones, to initiate infection. In contrast, anti-transcriptional protein had not been previously reported. The tail fiber protein and transporters, fimbriae chaperones were consistent with previous experimental discoveries, which indicated the robustness of our model and the effectiveness of model Interpretability. The novel factor, anti-transcriptional protein, showed the potential of model Interpretability. The feature weights with biological knowledge uncovered a largely unexplored phage-host interaction mechanism.

### Guidance for strain-level phage-host interacting prediction

Overall, the strategies to establish the precise and robust strain-level phage-host interaction prediction framework were summarized as follows: 1) The experimental dataset outperformed the manual dataset and the experimental negative can not be substituted. 2) Protein features had superior performance to DNA features. 3) For tailed phages, the combination of tail proteins and protein features for computational modeling can greatly reduce computation time and construction and were close to using whole-genome sequences in performance metrics.

## Discussion

In this study, we developed a unified framework to compare both the statistical architecture of phage–bacteria infection networks and the feature engineering strategies that enhance the performance of predictive models. Our analysis revealed that the infection network between *K. pneumoniae* strains and their associated bacteriophages exhibits a distinct modular structure with mild nestedness. In contrast, networks generated using the Leite method displayed a predominantly one-to-one interaction pattern, leading to a considerable number of falsenegative entries. These artificially introduced negatives significantly impair model performance, as true negative instances are essential for accurate learning particularly in imbalanced datasets.

Most studies have focused on identifying positive phage-host interactions. Due to the substantial cost and complexity of comprehensive phage-host characterization, experimentally validated antagonistic interactions (i.e., true negatives) remain largely absent from public datasets. Consequently, earlier efforts to model strain-level phage-host prediction have relied on strategies such as random sampling, clustering, or data augmentation to construct negative datasets. However, these synthetic approaches have seldom been evaluated rigorously owing to the lack of ground-truth negative data, often leading to overestimation of model performance and reduced generalizability.

Our dataset addresses this limitation by providing an unprecedented level of resolution and completeness, comprising nearly 13,000 experimental infection assays involving 125 phage isolates and 104 K. pneumoniae strains. Using this large-scale interaction matrix, we quantitatively demonstrated that negative datasets generated through random sampling introduce systematic biases that in-flate performance metrics (e.g. accuracy, precision, recall, AUPRC). Based on this finding, we recommend prioritizing experimentally validated datasets whenever possible for robust strain-level phage-host modeling.

Beyond dataset construction, we also evaluated the effectiveness of common feature extraction strategies. While both DNA-based (e.g., k-mer frequency) and protein-based (e.g., amino acid composition) features have been widely used for phage-host prediction, we found that protein-level features consistently outperformed DNA features across all tested scenarios. Notably, models using only tail protein sequences achieved performance comparable to those using wholegenome sequences, with the added benefit of reduced computational cost and faster model training. This result highlights the utility of targeting functionally relevant genomic regions, such as tail proteins, when designing efficient and accurate prediction pipelines.

Although our experimental dataset is currently limited to K. pneumoniae and its phages, the insights derived from it are broadly applicable. In particular, this dataset can serve as a benchmark for validating synthetic negative generation techniques or refining model optimization strategies in other host–phage systems. Given the prohibitive cost of large-scale experimental screening, leveraging available experimental negatives to inform more realistic modeling pipelines will be a critical direction for future research.

Another key feature of real-world phage-host datasets is the inherent class imbalance (e.g., a typical positive:negative ratio of ∼1:10), which poses challenges for traditional machine learning algorithms. Future work should explore imbalance-aware strategies such as custom loss functions (e.g., focal loss)[30] or generative adversarial networks (GANs) for data augmentatio[31]. As phages with highly specific infection profiles are increasingly recognized for their roles in disease modulation and microbiome dynamics, improving our understanding of phagehost specificity will significantly enhance microbial ecology and therapeutic research.

In conclusion, our work not only provides a high-quality experimental dataset but also delivers actionable guidance for selecting appropriate modeling strategies, thereby enabling more accurate and interpretable phage-host interaction predictions at the strain level.

## Methods

### Genotypic and phenotypic dataset collection

The genotypic dataset consisted of whole-genome sequences from 125 *K. pneumoniae* phages and 104 corresponding bacterial host strains. The phenotypic dataset included 13,000 experimentally validated phage-host interaction records. All data were obtained from the CNSA (CNGB Sequence Archive)[32] under accession number CNP0006217, available at: https://db.cngb.org/search/project/CNP0006217/.

### Bio-Community network analysis and mode detection

Phenotypic data were converted to the binary dataset, with 1 indicating positive infection and 0 indicating negative infection. 24 hosts were discarded as they showed full resistance to any of the phage (rows in the matrix with only 0). The *K. pneumoniae* phage–bacteria infection matrix with accompanying metadata and after cleaning can be found in the “Supplementary Data File 1.xlxs” file in the Supplementary Materials.

Modularity was quantified with the lpbrim[33] package in R using the findModules function with 1000 iterations, which indicated the presence of a dense cluster of related nodes embedded in the network. The Q_b_ value was used to measure the strength of the modularity of the bipartite network, where Q_b_ > 0.5 indicated a more pronounced modular structure. The Q_b_ was calculated as:

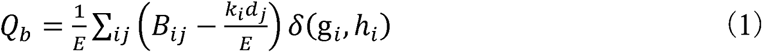

Where *B_ij_* represented the member of the bipartite matrix, indicating the presence or absence of a link between nodes *i* and *j*, i.e., a 1 or 0 in the matrix. *gi* and *h_i_* were the indexes of node *i* and *j,* and *k_i_*, *d_i_* was the degree of node *i, j E* represented the total number of links in the network.

Nestedness was measured with the oecosimu function from the vegan[34] package in R. The nestedness metric based on overlap and decreasing fill (NODF) index was originally proposed to evaluate the nestedness of binary networks. Nestedness was a typical feature of the mutualism network (plant–animal[35], plant–pollinator[36], mycorrhizal symbiosis[37], etc.), i.e. the mode of specialists from both sets of species interact preferentially with generalists. This NODF equation of binary dataset:

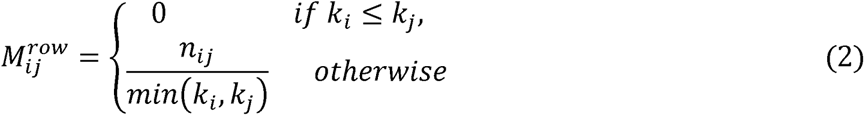

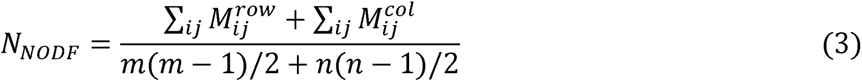

NODF measures the nestedness of matrices by computing their reduced filling and pairwise overlapping properties. NODF measures the nesting between rows by assigning a value)#$% for each row in the interaction matrix, where & is the number in row " and & is the number in row # number, !"*!" is the number of shared interactions between rows " and # (so-called pairwise overlap). Note that a positive contribution to NODF requires column pairs that satisfy the decreasing fill property, that is, when & > & is counted only if it is not counted, otherwise it counts as 0. Similarly)&$’ is used to calculate the column nesting contribution. The total nesting is the sum of the column and row nesting!" contributions.

### DNA and protein sequence feature calculation

DNA sequence feature was calculated by iLearn[38], including kmer, Reverse Compliment Kmer(RCKmer),Nucleic Acid Composition (NAC), Di-Nucleotide Composition (DNC), Tri-Nucleotide Composition (TNC), Composition of k-spaced Nucleic Acid Pairs (CKSNAP), Electron-ion interaction pseudopotentials of trinucleotide (PseEIIP), totally 64-dimensional vector.

Protein sequence feature was extracted from amino acid composition abundance, chemical element composition and molecular weight by calculating mean, maximum, minimum, standard deviation, variance, and median of each dimensional vector.

### Data diving and machine learning modeling

The binary phage-host interaction dataset was first preprocessed by encoding protein and DNA sequence features using amino acid frequency and k-mer frequency calculations, respectively. To ensure robust model evaluation, the dataset was randomly split into training (90%) and testing (10%) sets. This splitting was repeated 30 times with different random seeds to reduce sampling bias, and all results were averaged over these iterations.

We employed four supervised machine learning algorithms to build predictive models: Logistic Regression (LR)[39], Support Vector Machine (SVM)[40], Random Forest (RF)[41], and Extreme Gradient Boosting (XGBoost)[42]. These models were chosen to represent a spectrum of classification techniques, including linear models (LR), kernel-based methods (SVM), bagging ensembles (RF), and gradient boosting (XGBoost).

Hyperparameters for each model were optimized using grid search on the training data with 5-fold cross-validation. For LR and SVM, regularization strength and kernel parameters were tuned as 1.0 and rbf. RF parameters such as the number of trees and maximum depth were adjusted as 100 and None. XGBoost tuning focused on tree depth (6), learning rate (0.3), and subsampling ratios (1.0).

All analyses were implemented in Python (version 3.8.5) using the scikit-learn (v0.24.2) and XGBoost (v1.4.2) libraries. Computations were performed on a personal compution with Intel Xeon CPUs and 64 GB RAM. All code about modeling is available on github (https://github.com/a1678019300/PHIPrediction).

## Acknowledgements

This work is supported by National Key R&D Program of China (2020YFA0908700). We sincerely thank the China National GeneBank Data-Base (CNGB) for providing valuable data support and computational resources.

## Author Information

M.X. conceived the study. W.S. generated the raw data. M.L. and F.G compiled the training, validation, and test sets. L.M. and F.G implemented the algorithm. M.L., F.G and M.X. analyzed the results. M.L., F.G and M.X. drafted the manuscript and made the figures. M.X. and L.M. revised the manuscript. Jianqiang L. provided consultation. All authors read, edited, and approved the final manuscript.

## Supplementary Material

**Supplementary Figure 1.**
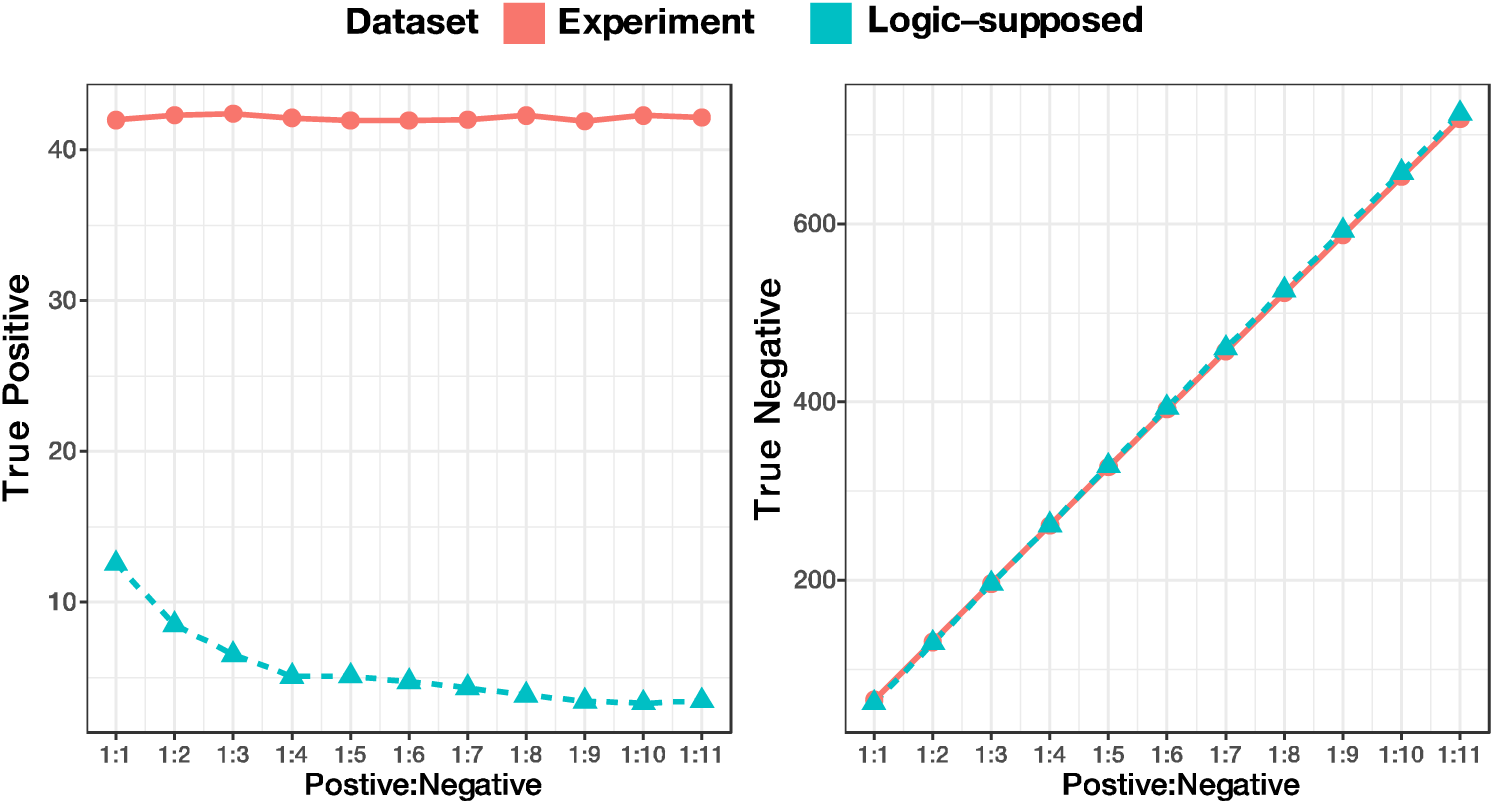
Effectiveness of Experimental and Random-Select Datasets on True Positive and True Negative.

**Supplementary Figure 2.**
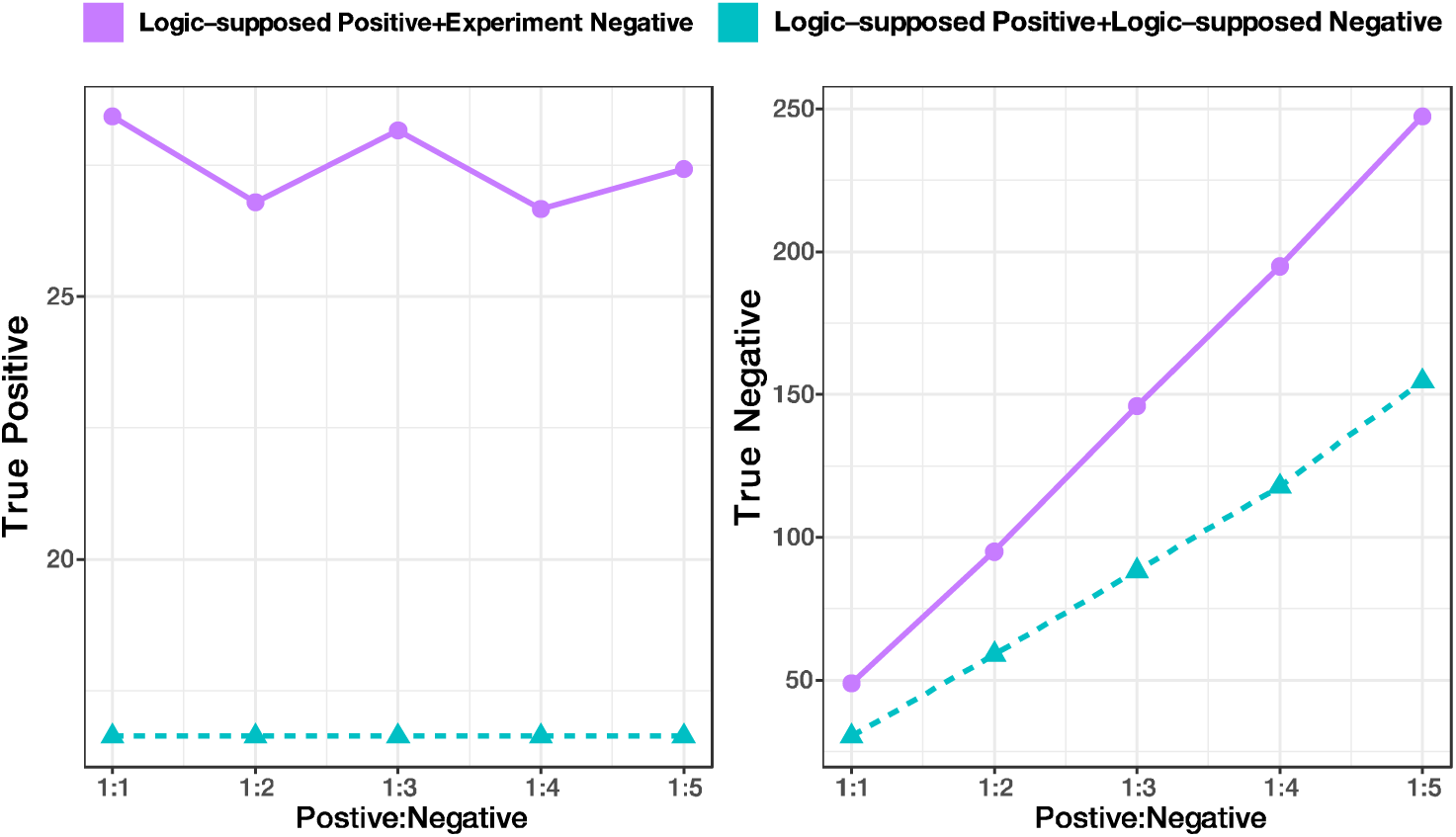
Effectiveness of Experimentally Derived Negative Data on True Positive and True Negative.

